# Data of the Insect Biome Atlas: a metabarcoding survey of the terrestrial arthropods of Sweden and Madagascar

**DOI:** 10.1101/2024.10.24.619818

**Authors:** A Miraldo, J Sundh, E Iwaszkiewicz-Eggebrecht, M Buczek, R Goodsell, H Johansson, BL Fisher, D Raharinjanahary, ET Rajoelison, C Ranaivo, C Randrianandrasana, J-J Rafanomezantsoa, L Manoharan, E Granqvist, LJA van Dijk, L Alberg, D Åhlén, M Aspebo, S Åström, A Bellviken, P-E Bergman, S Björklund, MP Björkman, J Deng, L Desborough, E Dolff, A Eliasson, H Elmquist, H Emanuelsson, R Erixon, L Fahlen, C Frogner, P Fürst, A Grabs, H Grudd, D Guasconi, M Gunnarsson, S Häggqvist, A Hed, E Hörnström, H Johansson, A Jönsson, S Kanerot, A Karlsson, D Karlsson, M Klinth, T Kraft, R Lahti, M Larsson, H Lernefalk, Y Lestander, L-T Lindholm, M Lindholm, U Ljung, K Ljung, J Lundberg, E Lundin, M Malmenius, D Marquina, J Martinelli, L Mertz, J Nilsson, A Patchett, N Persson, J Persson, M Prus-Frankowska, E Regazzoni, K-G Rosander, M Rydgård, C Sandblom, J Skord, T Stålhandske, F Svensson, S Szpryngiel, K Tajani, M Tyboni, C Ugarph, L Vestermark, J Vilhelmsson, N Wahlgren, A Wass, P Wetterstrand, P Łukasik, AJM Tack, AF Andersson, T Roslin, F Ronquist

## Abstract

We present the data from the Insect Biome Atlas project (IBA), characterizing the terrestrial arthropod faunas of Sweden and Madagascar. Over 12 months, weekly Malaise trap samples were collected at 203 locations within 100 sites in Sweden and at 50 locations within 33 sites in Madagascar; this was complemented by soil and litter samples from each site. The field samples comprise 4,749 Malaise trap, 192 soil and 192 litter samples from Sweden and 2,566 Malaise trap and 190 litter samples from Madagascar. Samples were processed using mild lysis or homogenization, followed by DNA metabarcoding of COI (418 bp). The data comprise 698,378 non-chimeric sequence variants from Sweden and 687,866 from Madagascar, representing 33,989 (33,046 Arthropoda) and 77,599 (77,380 Arthropoda) operational taxonomic units, respectively. These are the most comprehensive data presented on these faunas so far, allowing unique analyses of the size, composition, spatial turnover and seasonal dynamics of the sampled communities. They also provide an invaluable baseline against which to gauge future changes.

## Background & Summary

Insects comprise the vast majority of macroscopic biodiversity ^1^ and are crucial for ecosystem functioning ^2–5^. Yet, despite their ecological importance and remarkable diversity, insects remain one of the least understood organism groups. An estimated 70-85% of insect species still remain unknown to science ^6,7^ and for the large majority of those described, we know little about their biology, distribution, or population trends ^8^.

Recent studies have indicated a decline in terrestrial insect biomass and diversity during the last decades ^9–16^, but the data are often limited in taxonomic, geographic and/or temporal scope, hindering comprehensive conclusions to be reached ^17–21^. It is clear that existing data on insect biodiversity provide only fragmented insights into how many insect species there are, how they are structured into communities and how these communities contribute to healthy ecosystems.

Comprehensive longitudinal sampling across and within regions is urgently needed to close this knowledge gap. However, setting up such long-term monitoring projects faces many challenges, from financial constraints, complex management and coordination hurdles to methodological limitations in the processing and identification of collected material. Indeed, a significant barrier has been the absence of cost-effective methods for the comprehensive inventorying of insect faunas. For example, one of the most ambitious countrywide insect inventory projects to date, the Swedish Malaise Trap Project (SMTP), collected an estimated 20 million specimens at 50 sites over a three-year period ^22^. Sorting the material into 350 taxonomic fractions suitable for specialist processing took 15 years. During this time period, only 0.85% of the specimens had been identified to species, despite a substantial effort from numerous taxonomists ^22,23^.

Fortunately, in recent years, the traditional insect inventorying methods relying on morphological identification by taxonomic experts have been complemented by considerably faster and more cost-efficient methods for processing and identifying insect samples, relying on DNA metabarcoding. This has revolutionized our capacity to comprehensively study and effectively monitor insect faunas ^24^.

In the Insect Biome Atlas (www.insectbiomeatlas.org) project, we used DNA metabarcoding of an extensive set of community samples to characterize the terrestrial arthropod faunas of Sweden and Madagascar, and their associated microbiomes, in unprecedented detail. Sweden and Madagascar were chosen because they represent two countries of approximately the same size but with completely different biogeographic and geological history. The Swedish fauna has been assembled by independent migration from different refugia after the last glacial maximum, some 20,000 years ago ^25^. In contrast, the Madagascar fauna has largely evolved in place, and in isolation ^26^. The environmental factors also contrast sharply: Sweden is characterized by a northern temperate climate and an elevational gradient running mainly from the east coast to the Scandinavian mountains in the west, whereas Madagascar is marked by a tropical climate and a significantly more complex topographical setting. Thus, any similarities between Sweden and Madagascar clearly represent non-random structuring of the faunas, while the differences between them illustrate much of the range of global variation one might expect among faunas.

Over a period of 12 months in 2019-20, weekly Malaise trap samples were collected at 100 sites (203 locations) in Sweden and 33 sites (50 locations) in Madagascar (Figure 1). Sites and trap locations were selected to provide a representative sample of Swedish habitat types. In brief, sites were stratified across the habitats in rough proportion to the relative coverage of habitats in the Swedish landscape ^27^. In Madagascar, only forests (National Parks) were targeted due to logistical constraints. A hierarchical design involving both single-trap and multitrap sites was used to allow more powerful analyses of spatial turnover. The Malaise traps were maintained by citizen science volunteers in Sweden and park rangers in Madagascar. This approach not only substantially mitigated sampling-related challenges but also engaged a diverse range of stakeholders in the scientific process.

**Figure 1.**
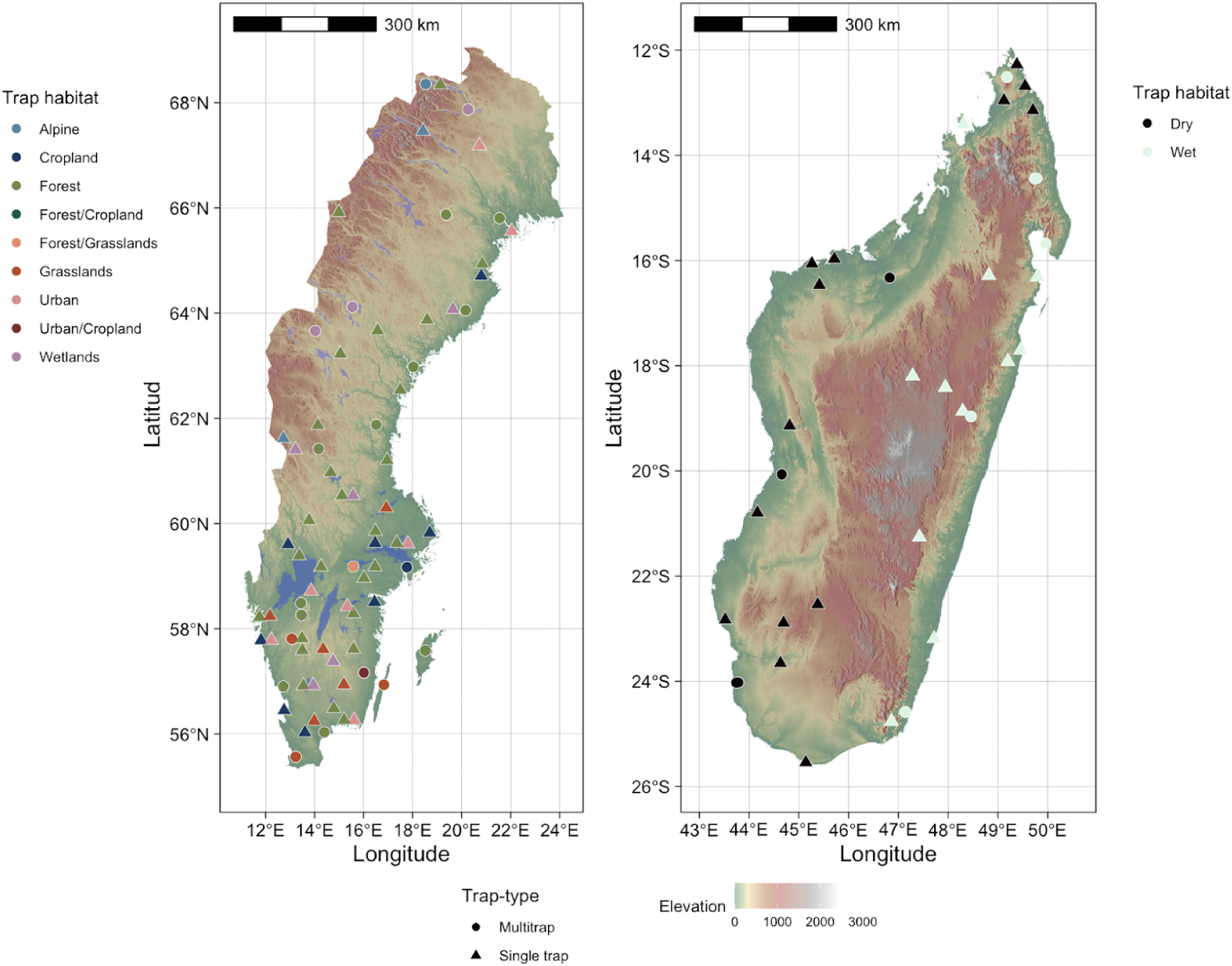
Sampling sites for Sweden (left) and Madagascar (right). Sampling sites with multiple Malaise traps (multitrap sites) are depicted with filled circles and single trap sites with filled triangles. Colours of each point denote the habitat the trap was placed in. In Sweden (left panel) a total of 203 Malaise traps were placed across 100 sampling sites: 102 traps placed in forests, 33 in croplands, 26 in wetlands,16 in grasslands, 14 in alpine and 12 in urban areas for Sweden. In Madagascar (right panel), sampling sites were associated with forested habitats located within protected areas only. Fifty Malaise traps across 33 sites were deployed: 27 Malaise traps in rainforests (16 sites, denoted in white) and 23 Malaise traps in dry forests (17 sites, denoted in black). Background map colour illustrates elevation.

To further characterize soil arthropods and soil microbiomes, we complemented the Malaise trap samples with soil and litter samples. In addition, we collected environmental data and data on ecosystem functions at each site. In total, the project yielded 4,753 Malaise trap samples, 188 soil samples and 189 leaf litter samples from Sweden and 2,566 Malaise trap samples and 190 leaf litter samples from Madagascar.

Samples were processed using mild lysis or homogenization, followed by DNA metabarcoding of COI (418 bp) on the NovaSeq6000 platform using optimized protocols developed in the project ^28–30^. A novel pipeline for processing the metabarcoding data - developed in the project - was used for quality filtering and taxonomic characterization of the samples based on these data ^31^. This pipeline utilizes the extensive amounts of samples collected in the project to provide data of significantly improved quality and accuracy compared to the current state of the art.

Here we present the key datasets resulting from the project. The data on ecosystem functioning and arthropod biomass have been published elsewhere ^32,33^ and the data on soil microbiome (bacterial 16S and fungal ITS2) and microbiome associated with the arthropod communities in Sweden will be presented separately.

The COI data presented here comprise 698,378 non-chimeric sequence variants from Sweden and 687,866 from Madagascar, representing 33,989 and 77,599 operational taxonomic units (OTUs), respectively, after filtering and cleaning. The Swedish OTUs were distributed across 11 phyla, 26 classes and 97 orders. Of the OTUs, 79%, 50%, 34% and 60% were classified at the level of Family, Genus, Species and BOLD BIN ^34^, respectively. The low annotation success at the species level compared to the BOLD BIN level is partly due to name resolution problems for BOLD BINs—such as several synonyms being in active use, or the same species being placed in different genera by different annotators—and partly due to failures of CO1 barcodes in accurately circumscribing closely related species. Distinguishing between these cases would require detailed analysis on a case-by-case basis. For Madagascar, the OTUs were distributed across 9 phyla, 21 classes and 90 orders, with 28%, 4%, 2% and 9% of OTUs classified at the level of Family, Genus, Species and BOLD BIN, respectively. For both datasets, the phylum Arthropoda accounted for ≥99.5% of total reads and >97% of OTUs. Calibration curves relating biomass to the number of specimens suggest that the analyzed Malaise trap samples comprise around 7.0 million insects from Sweden and 1.7 million insects from Madagascar.

The datasets we present here offer profound insights into the diversity, composition and structure of the terrestrial arthropod communities of Sweden and Madagascar. They also provide an invaluable baseline against which to gauge future changes in these faunas. While the Insect Biome Atlas project focuses exclusively on Sweden and Madagascar, we expect that it will serve as a template for similar studies in other regions. We also hope that it encourages a wider interest in comprehensive studies of insect faunas and in establishing long-term monitoring of insect populations globally.

## Methods

### Site selection

In Sweden, sampling sites were selected following a stratified design based on the major landscape types as identified by the National Inventory of Landscapes in Sweden - NILS ^27^. A total of 203 Malaise traps were deployed within six major habitats across 100 sites: 102 traps in forests, 33 in croplands, 26 in wetlands,16 in grasslands, 14 in alpine and 12 in urban areas (Figure 1). The proportion of Malaise traps by habitat is representative of the contribution of each habitat type to the overall Swedish landscape (Table 1), with down-weighing of the most common habitat (forests, that account for 60% of the landscape according to NILS) and some up-weighing of the remaining, sparser habitats (croplands and wetlands, which account for 8% of the landscape each; and urban sites and grasslands, which each account for 3% of the landscape). At 21 of the 100 sampling sites, we implemented a hierarchical design by deploying up to 6 Malaise traps instead of one. In Sweden there were 12 multi-trap sites with between 2 and 6 traps, and in Madagascar there were 9 multi-trap sites with between 2-4 traps.

**Table 1.** Number (n) and proportional representation (%) of Malaise traps per habitat and area (m2) and proportion (%) of each habitat in Sweden. Data on area of each habitat was retrieved from the National Inventory of Landscapes in Sweden ^27^. Note that area presented here does not correspond to the total area of habitats present in sweden but only to the sampled habitat plots in NILS.

| **Habitat** | **Area** |  | **Malaise traps** |  |
| --- | --- | --- | --- | --- |
|  | **m<sup>2</sup>** | **%** | **n** | **%** |
| **alpine** | 170195 | 8 | 14 | 7 |
| **aquatic** | 205368 | 9 | 0 | 0 |
| **croplands** | 163551 | 8 | 33 | 16 |
| **forest** | 1294787 | 60 | 102 | 50 |
| **grasslands** | 64772 | 3 | 16 | 8 |
| **substrate** | 14430 | 1 | 0 | 0 |
| **urban** | 74943 | 3 | 12 | 6 |
| **wetlands** | 177123 | 8 | 26 | 13 |
| **Total** | 2165170 | 100 | 203 | 100 |

Distances between Malaise traps at these multitrap sites ranged from 90 to 1200 meters, sampling two different habitats when possible (three Malaise traps per habitat). In Madagascar, sampling sites were associated with forested habitats located within protected areas only. In total we deployed 50 Malaise traps across 33 sites: 27 Malaise traps in rainforests (16 sites) and 23 Malaise traps in dry forests (17 sites) (Fig 1). At eight of the 33 sites, we implemented a hierarchical design by deploying 3 traps within a single habitat. Distances between Malaise traps at these multitrap sites ranged from 100 to 4700 meters. Malaise traps were set up by a team of fieldworkers together with the trap managers at each site: 150 volunteers in Sweden and 107 local community members and park rangers in Madagascar. At the time of setting up the Malaise traps, trap managers were trained on how to set up and maintain the trap, retrieve the samples from the trap and collect metadata associated with the sampling event.

### Standing characteristics and other abiotic data

At each Malaise trap location in Madagascar we measured a set of standing characteristics. To assess tree density we set up four 10 m long transects, one on each side of the trap, and measured the diameter at breast height (DBH) of all trees with its roots or stem within 10 cm of the transect. For multi-stemmed trees, we measured only the DBH of the largest stem. We included measurements for trees, lianas (L), palms (P) and ferns (F). We also measured the circumference of all trees with a DBH larger than 30 cm within a 10 m radius of the Malaise trap. Finally we measured canopy cover at each Malaise trap location by taking five photographs of the canopy at 2 meters height straight up to the canopy. One photograph was taken at the center of the Malaise trap and the other four photographs were taken five meters away from the trap, one on each side of the trap. Each photograph was analysed with ImageJ software to get a percentage of canopy cover.

At each Malaise trap location for both Sweden and Madagascar we also measured soil nutrients and soil pH. Topsoil (0-20cm) was sampled at 5 sites around each Malaise trap: one soil core (6 cm diameter) at the center of the trap and one soil core on each of the four “sides” of the trap, five meters away from the trap. Soil samples were taken as composite samples from the five locations. At each site, prior to taking the soil cores, we measured leaf litter depth using a ruler and soil humidity with the SM150 soil moisture kit. Soil samples from Sweden were sent to Eurofins to measure the following macronutrients: AL-extractable P, K, Mg and Ca; NH_4_; NO_3_; K/Mg ratio and mineral N (Nmin). The concentrations of soil nutrients are presented as mg·100 g^−1^ air-dry sieved soil (<2 mm). Soil samples from Madagascar were sent to the Laboratoire des Radioisotopes to measure the following macronutrients: organic C (g/Kg), total N (g/Kg), total P (g/Kg), exchangeable K (cmol/Kg), exchangeable Ca (cmol/Kg) and exchangeable Mg (cmol/Kg).

### Sampling for biotic characterization

#### Malaise trap samples

Arthropods were collected in individually barcoded bottles pre-filled with 400 mL of 95% ethanol attached to each Malaise trap. To facilitate the recording of metadata associated with each sampling event, trap managers used a mobile application specifically designed for the project. The app allowed scanning the individually barcoded Malaise trap and sample bottle at the time of collection so that each sampling event was automatically associated with a specific trap. When the barcode of each sample was scanned, the app automatically registered the GPS coordinates and the date and time of the collection event in addition to any other metadata manually inserted by the user, such as the trap condition at collection. In Sweden, Malaise traps were active between January and December 2019. Samples were collected every week during spring and summer (March/April to September/October depending on latitude) and monthly or bi-weekly in the autumn and winter (October/November to March/April, depending on latitude), resulting in 4,753 insect community samples. In Madagascar, Malaise traps were active between August 2019 and July 2020. Samples were collected every week throughout that period, resulting in 2,566 insect community samples. Every fourth sample from each trap location (638 samples in total, roughly one sample per month), was left at the Madagascar Biodiversity Center (https://www.madagascarbio.org/) to help build a natural history collection of Malagasy insects in the country of origin. The remaining samples were shipped to Sweden for DNA extraction and metabarcoding as described below.

#### Soil and litter samples

Soil and leaf litter arthropod communities were sampled during the growing season (26th of June 2019 to 27th of July 2019) at each Malaise trap location in Sweden. Leaf litter arthropod communities were sampled by collecting a total volume of 1 L of leaf litter from five different sites around each Malaise trap, 2.5 meters away from the nearest extremity of the trap. Soil arthropod communities were sampled using soil cores (5.5 cm diameter) with a depth of 10 cm at exactly the same five sites where litter samples were taken. The five soil core samples from each Malaise trap location were pooled and mixed by hand immediately after collection, then stored in a cool place. Within 48h after collection, living arthropods were extracted from soil and litter samples over a period of 96 hours using funnel extractors ^35^. To cause the arthropods to exit the substrate before being trapped in the dried-out structure, we gradually increased the temperature over the first 24 hours to a temperature of 52 °C. This temperature was then held constant for the remaining 72 hours. Arthropods were collected directly into 100 mL plastic bottles filled with 95% ethanol. Leaf litter arthropod communities were also sampled at each Malaise trap location in Madagascar. Here we collected four samples in each direction of the Malaise trap (back, front, left and right), five meters away from it. Leaf litter sampling involved sifting and concentrating 2 L of leaf litter from a 1 m^2^ square using a Winkler-sifter. Before sifting, the leaf litter was minced with a machete to dislodge any insects hiding in the twigs or decayed logs. After sifting, the leaf litter was stored in a cloth bag before extracting the arthropods. Within 12 hours of sampling, living arthropods were extracted at ambient temperature overnight into a 100 mL bottle filled with 95% ethanol using a mini-Winkler extractor for a total of 12 hours ^36^.

### DNA extraction

#### Malaise trap samples

DNA was extracted from Malaise trap samples using both non-destructive (mild lysis and preservative ethanol) and destructive methods (homogenization) (Figure 2).

1. Mild lysis: DNA was extracted from 6,483 Malaise trap samples (4,560 from Sweden and 1,923 from Madagascar) using a non-destructive mild-lysis protocol (FAVIS protocol, steps 1-17 ^29^). In brief, ethanol was first decanted from each sample and insect biomass was measured. Lysis buffer, proteinase K and biological spike-ins (specimens from foreign species, ie, species that don’t occur in the respective countries) were added and samples were incubated for 2h45m at 56°C in a dry shaking incubator. After the incubation period the lysate was drained and each insect community was remixed with the previously-decanted ethanol for long-term storage. DNA was purified from 225 μL of lysate using silica-coated magnetic beads with the KingFisher Cell and Tissue DNA kit on a KingFisher Flex 96 robot (both Thermo Scientific) according to manufacturer instructions. The list and number of biological spike-ins added in each country (SE and MG) can be retrieved at https://doi.org/10.17044/scilifelab.25480681.v537.
2. Homogenization: Approximately every fourth sample collected at each sampling site in Sweden (n=873) was further processed using a destructive homogenization protocol ^30^ after mild lysis. In brief, after decanting preservative ethanol we homogenized each bulk sample into an “arthropod soup” using the Ultra Turrax Drive homogenizer (IKA™). After homogenization, the entire “arthropod soup” was digested with lysis buffer and proteinase K for 2h45min at 56°C in a dry shaking incubator. To make sure that we used all available DNA from each sample, we combined the homogenate with the respective lysate obtained during mild lysis before proceeding with DNA purification. At this time we also added a standardized amount (5 million copies) of two synthetic oligonucleotide sequences (synthetic spike-ins) to each homogenate aliquot. Synthetic spike-ins were produced as described in et al. ^38^ and their sequences can be found at https://doi.org/10.17044/scilifelab.25480681.v5 ^37^. DNA was purified from a 225 µL subsample of homogenate using silica-coated magnetic beads with the KingFisher Cell and Tissue DNA kit on a KingFisher Flex 96 robot (both Thermo Scientific) according to manufacturer instructions.
3. Preservative ethanol: For 15 Malaise trap samples we also extracted DNA from the preservative ethanol (prior to mild lysis and homogenization). Ethanol was first decanted from each sample as described in step 7 of the FAVIS protocol ^29^. The decanted ethanol was then manually filtered using a Millipore® Sterivex™ filter unit (pore size of 0.22 μm). After filtration, DNA was lysed inside the Sterivex unit by adding 540 µl of lysis buffer (ATL, Qiagen) and 60 µl proteinase K (Qiagen) and incubating the unit at 56°C overnight. DNA was transferred from the Sterivex unit into a DNeasy Blood and Tissue Kit column (Qiagen) for purification, following manufacturer instructions.

**Figure 2.**
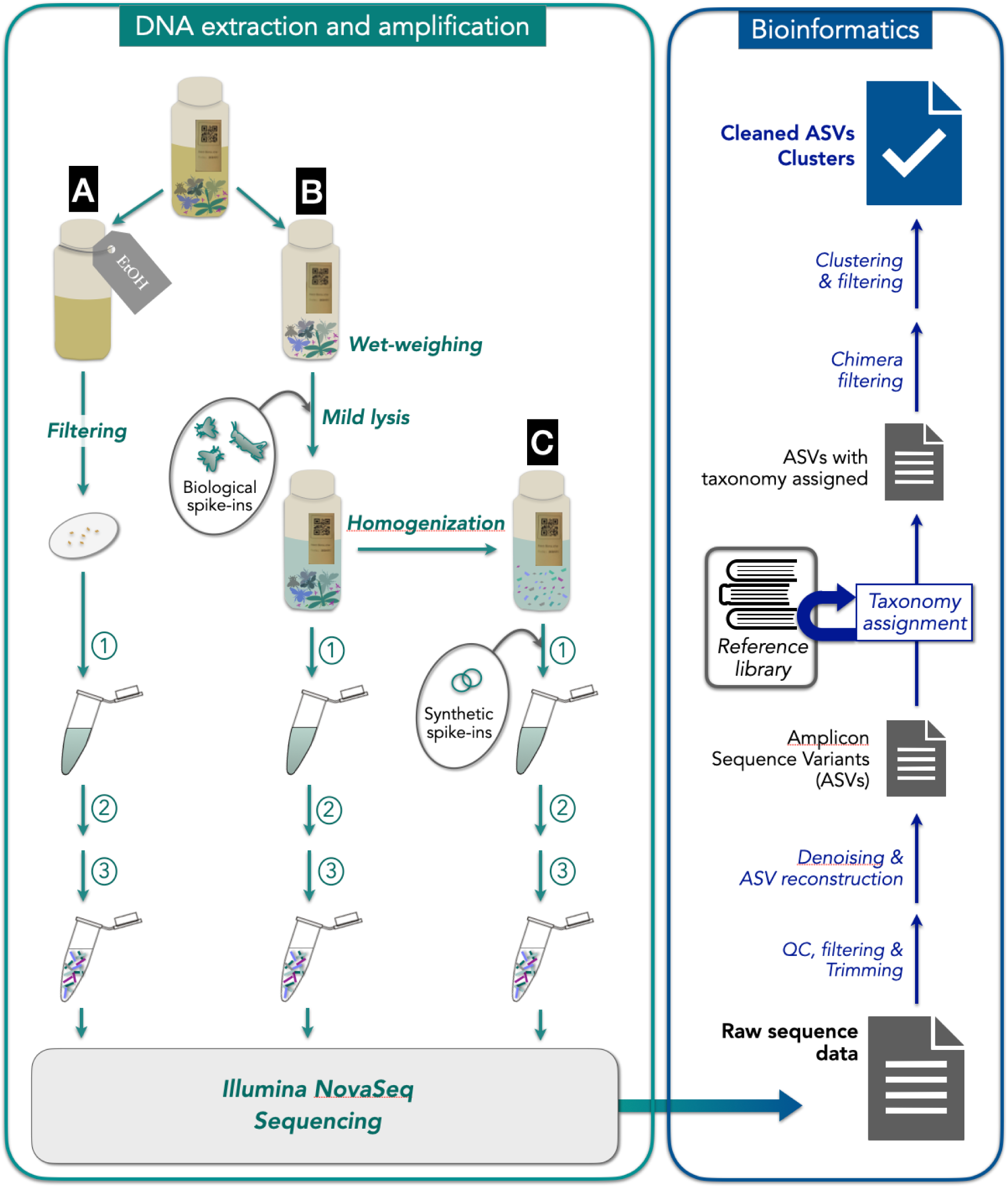
**Left panel**: Malaise trap sample processing in the lab. DNA was extracted from (A) the ethanol filtered from malaise trap samples; (B) the lysates using the FAVIS mild lysis protocol ^29^; and (C) the homogenates ^30^. Before lysis, biological spike-ins and synthetic spikes were added to the samples in B and C, respectively. After lysis, DNA was purified using silica-coated magnetic beads with the KingFisher Cell and Tissue DNA kit on a KingFisher Flex 96 robot (1). After DNA purification, a 418 bp fragment of the mtDNA cytochrome c oxidase subunit 1 (COI) gene was amplified using a 2-step PCR approach. In the first step (2), the target region was amplified using broad-spectrum primers BF3 *CCHGAYATRGCHTTYCCHCG* ^42^ and BR2 *CDGGRTGNCCRAARAAYCA* ^43^. In the second step (3), indexes were added and Illumina adapters completed. Samples were pooled and library pools were sequenced on a NovaSeq6000 instrument using the ‘NovaSeqXp’ workflow in ‘SP 500-cycle’ flow cells, with 384 double-uniquely indexed samples per lane. **Right panel**: schematic representation of processing sequence data from raw reads to ASV clusters. Briefly, paired end reads were trimmed in a series of steps using the program cutadapt ^47^ and filtered to retain only sequences that were between 403 and 418 nt in length (in incremental steps of 3 nt) and that did not contain any in-frame stop codons. Preprocessed reads were then denoised using DADA2 ^49^ to infer amplicon sequence variants (ASVs). The ASVs were then processed using the HAPP pipeline ^31^ to taxonomically annotate the ASVs (using SINTAX classifier and a purposely built CO1 database). Taxonomic assignments obtained from SINTAX were refined using phylogenetic methods with EPA-NG ^55^ into an insect phylogeny ^56^, followed by taxonomic assignment with gappa ^57^. ASVs were then clustered into OTUs using Swarm (ref). Finally, the clustered data was cleaned from NUMTs and other types of noise using the NEEAT algorithm ^31^.

#### Soil and litter samples

Arthropod litter samples from Madagascar were processed using the same mild lysis protocol used for the Malaise samples described above, with the exception of not adding biological spike-ins to the samples. For Sweden, we used a 1mm wire-sieve to separate each sample (from soil or litter) into two components based on size: the macrofauna subsample, composed mainly of adult and larval Coleoptera, and the mesofauna subsample, dominated by mites and springtails. To remove debris and dirt accumulated in the mesofauna subsamples, we processed them further using a combination of flotation in distilled water, adapted from the Flotation-Berlese-Floatation protocol in ^39,40^, followed by vacuum pump filtration (41 µm nylon filter). DNA was extracted separately from the macrofauna and the mesofauna subsamples using the Kingfisher Cell and Tissue DNA kit. Ethanol was first dried from each subsample in a dry incubator at 40°C for 4-5 hours. After drying, specimens were manually homogenized with a pestle and lysed overnight at 56°C by adding 800 µL of lysis buffer and 100 µL of proteinase K (provided in the Kingfisher Cell and Tissue DNA kit). DNA was then purified from a 225 µL aliquot of homogenate using silica-coated magnetic beads with the KingFisher Cell and Tissue DNA kit on a KingFisher Flex 96 robot (both Thermo Scientific) following manufacturer instructions. DNA extracts were quantified using Qubit™ dsDNA HS Assay Kit on a Qubit Fluorometer (INVITROGEN™), and the original sample was reassembled by combining DNA extracts from each subsample pair in a ratio of 1:10 (amount of DNA of macrofauna: amount of DNA of mesofauna). This minimizes the bias in the sequencing depth due to biomass differences between the two components of each sample ^39,40^.

### Library preparation and sequencing

To characterize arthropod communities, we amplified a 418 bp fragment of the mtDNA cytochrome *c* oxidase subunit 1 (COI) gene, using the purified DNA extracted from the Malaise trap samples (lysates, homogenates and preservative ethanol) and soil and litter samples. Each bulk sample was metabarcoded using a two-step PCR approach for library preparation (method 4 of ^41^). In the first step (PCR 1), the target region was amplified using broad-spectrum primers BF3 *CCHGAYATRGCHTTYCCHCG* ^42^ and BR2 *CDGGRTGNCCRAARAAYCA* ^43^. The primers were supplemented with 5′-end Illumina sequence adapters (forward: ACACTCTTTCCCTACACGACGCTCTTCCGATCT-3′, reverse: 5′-GTGACTGGAGTTCAGACGTGTGCTCTTCCGATCT). To increase the complexity of the libraries, each primer was further complemented with variable length inserts (TGA, GA, A, or no base for the forward primer, and GAT, AT, T, or no base for the reverse primer)between the adapter sequence and the target-binding region, generating phased primers in equal proportions ^44,45^. PCR 1 reactions were carried out in a final reaction volume of 40 μL containing 20 μL QIAGEN Multiplex PCR Master mix, 1 μM of each primer, and 4 μL of template DNA (for mild lysis and preservative ethanol samples). As the DNA concentration for homogenates was relatively higher, the PCR reactions were carried out in a final volume of 10 μL with 1 μL of template DNA. The PCR conditions were 95 °C for 15 min, 25 cycles of 94 °C for 30 s, 50 °C for 90 s and 72 °C for 90 s, followed by a final elongation step of 72 °C for 10 min. PCR 1 products were cleaned with magnetic beads (Carboxyl-modified Sera-Mag Magnetic Speed-Beads, Hydrophobic, CYTIVA), using 2/1 (v/v) magnetic beads to sample ratio. In the second step (PCR 2, indexing PCR), indexes were added and Illumina adapters completed. The indexing primer design followed the Adapterama scheme ^41,46^ with 10 bp indexes as follows: index i5 (forward): AATGATACGGCGACCACCGAGATCTACACxxxxxxxxxxACACTCTTTCCCTAC index i7 (reverse): CAAGCAGAAGACGGCATACGAGATxxxxxxxxxxGTGACTGGAGTTCAG. Libraries were double-uniquely indexed - in other words, each forward and each reverse index was used for only one library in a given sequencing lane. PCR 2 conditions were 95°C for 15 min, 7 cycles of 94 °C for 30 s, 50 °C for 90 s and 72 °C for 90 s, followed by a final elongation step of 72 °C for 10 min. To form the sequencing pool, all samples were pooled approximately equimolar based on the intensity of the band in an agarose gel, and the resulting sample pool was then purified with the Promega ProNex® Size-Selective Purification System, using 1/1.5 (v/v) pool to magnetic beads ratio. The quality of the library pool was then checked with an Agilent DNA High Sensitivity Kit on a 2100 Bioanalyzer instrument (Agilent Technologies). Library pools were sequenced on a NovaSeq6000 instrument using the ‘NovaSeqXp’ workflow in ‘SP 500-cycle’ flow cells, with 384 double-uniquely indexed samples per lane, or a total of 768 libraries per flow cell (8×96 well plates). This sequencing was performed at the Swedish National Genomics Infrastructure (NGI) at SciLifeLab (Solna, Sweden). The detailed step-by-step protocol can be found in ^29^.

### Processing sequencing data

Sequences were preprocessed (read trimming and filtering) using a Snakemake workflow available at <u>github.com/biodiversitydata-se/amplicon-multi-cutadapt</u>. In this workflow, paired end reads were trimmed in a series of steps using the program cutadapt ^47^ (v3.1):

1. Discard all reads with the Illumina TruSeq adapters in either the 5’ or 3’ end of sequences.
2. Search for and trim primer sequences from the start of reads in R1 and R2 files using forward and reverse primers, respectively. Remove any untrimmed reads. This step is done with additional settings ‘--no-indels’ and ‘-e 0’ in order to only accept perfect matches.
3. Discard any remaining reads that still contain primer sequences.
4. Trim reads to a fixed length. This length is calculated by subtracting the length of the longest primer from the read length defined by the ‘expected_read_length’ parameter under the cutadapt: section in the config file (default value is 251).

Reads were then filtered to retain only sequences that were between 403 and 418 nt in length (in incremental steps of 3 nt) and that did not contain any in-frame stop codons. Preprocessed reads were denoised using the <u>nf-core/ampliseq</u> Nextflow workflow ^48^ (v2.4.0) which uses the DADA2 algorithm ^49^ to infer amplicon sequence variants (ASVs) from the preprocessed reads. Due to the indexing scheme used with unique dual indexes per sample it was not necessary to perform per-sample abundance filtering to correct for mistagging ^50,51^.

The ASVs were then processed using the HAPP pipeline (https://github.com/insect-biome-atlas/happ) described separately ^31^ to taxonomically annotate the ASVs, remove chimeras, cluster the ASVs into OTUs, and remove NUMTs and other noise from the data. Specifically, ASVs were taxonomically annotated using SINTAX ^52^, as implemented in vsearch ^53^, against a custom made reference COI database available at https://doi.org/10.17044/scilifelab.20514192.v4 ^54^. This reference database was assembled from sequences in the BOLD database ^34^ as follows. Firstly, sequence and taxonomic information for records in BOLD were downloaded from the <u>GBIF Hosted Datasets</u>. This data was then filtered to only keep records annotated as ‘COI-5P’ and assigned to a BOLD BIN ID. The taxonomic information was parsed in order to assign species names and resolve higher-level ranks for each BOLD BIN ID. Sequences were processed to remove gap characters and leading and trailing ‘N’s.

After this, any sequences with remaining non-standard characters were removed. Sequences were then clustered at 100% identity using vsearch. This clustering was done separately for sequences assigned to each BOLD BIN ID. These steps were implemented in a Python package called coidb, available at https://github.com/insect-biome-atlas/coidb. Taxonomic assignments obtained from the kmer-based SINTAX classifier were refined using phylogenetic methods. Specifically, ASV sequences classified as Insecta or Collembola (class) but unclassified at order level by SINTAX were reassigned by phylogenetic placement with EPA-NG ^55^ into an insect phylogeny ^56^, followed by taxonomic assignment with gappa ^57^. Assignments obtained this way were used to update the SINTAX taxonomy, but only at the order level, leaving child ranks with the ‘unclassified’ prefix.

Following taxonomic assignments ASVs were further processed to remove chimeras with uchime ^58^ and the remaining non-chimeric sequences were clustered using Swarm ^59^ (v3.1.0) with setting ‘-d 15’. Taxonomic assignments for ASVs within clusters were resolved to the cluster level by an abundance-based consensus approach. For each cluster, starting from the lowest rank (BOLD BIN here), each unique taxonomic name was weighted by the sum of reads across samples and ASVs with said name. These sums were then normalized to percentages. If a single taxonomic name made up at least 80%, then that name was assigned to the cluster, including the assignments of parent ranks. If no single name reached the 80% consensus threshold, the process was iterated for the parent rank. Ranks for which no consensus could be reached were prefixed with ‘unresolved.’ followed by the name of the most resolved consensus taxonomy. For each cluster, the ASV sequence with the highest median of normalized read counts across samples was selected as representatives of the cluster. Ties were broken by taking the ASV with the highest mean.

The clustered data was further cleaned from NUMTs and other types of noise using the NEEAT algorithm, which takes taxonomic annotation, correlations in occurrence across samples (‘echo signal’) and evolutionary signatures into account, as well as cluster abundance ^31^. We used default settings for all parameters in the evolutionary and distributional filtering steps, and removed clusters unassigned at the order level and with less than three reads summed across each dataset. Additionally we removed clusters present in more than 5% of blanks.

### Biomass and count data

To allow an assessment of how the biomass of a Malaise trap sample translates to the number of specimens, we counted all the specimens for 24 Malaise trap samples from Sweden. We complemented these data with biomass estimates and specimen counts for 224 Malaise trap samples from a separate Swedish Malaise trapping campaign in 2018–2019, the Swedish Insect Inventory Project (SIIP).

The data (Figure 3) show that the number of specimens is only approximately proportional to the biomass of a sample (linear model without intercept, adjusted R^2^ 0.81; Figure 3a). Specifically, there is a slight tendency for the larger samples (in terms of biomass) to contain more specimens than if the relation was strictly proportional, as shown in a log-log model (R^2^ 0.79, regression coefficient 1.09 ± 0.04; Figure 3b). Fitting a linear model with intercept does not support the alternative explanation of a constant amount of alcohol residue causing this (adjusted R^2^ 0.69, intercept positive and not negative as expected under the hypothesis). Using the proportional model, the Swedish Malaise trap material is estimated to contain 7.0 M specimens, and the Madagascar material 1.7 M specimens. Accounting for the deviation from proportionality by applying the log-log regression equation to each sample separately, the Swedish material is instead estimated to contain 5.6 M specimens and the Madagascar material 1.2 M specimens.

**Figure 3.**
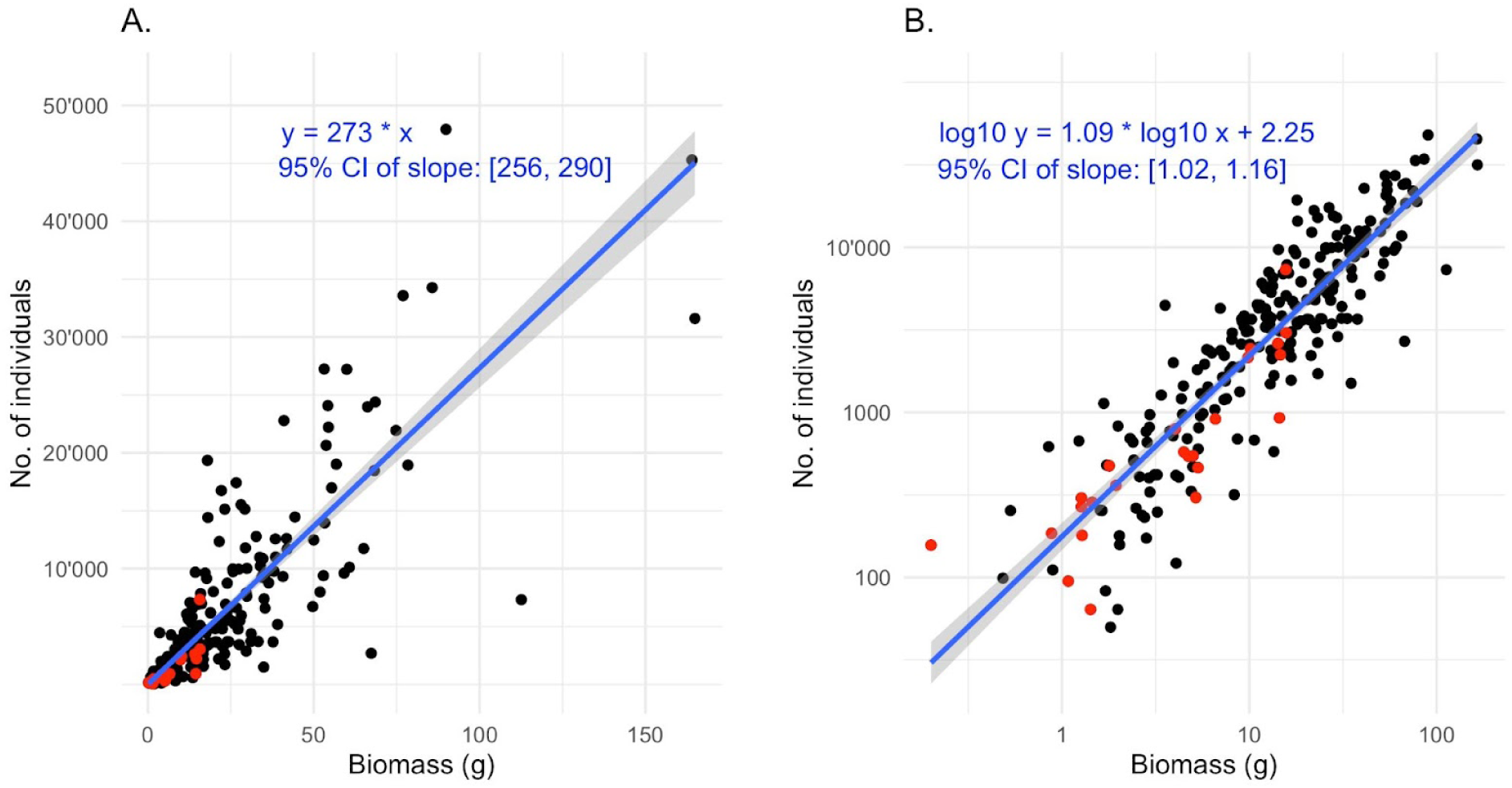
Linear regression between biomass of Malaise trap samples and the number of individuals (insect specimens). The data points are individual Malaise trap samples; the 224 black data points are from the SIIP project and the 24 red data points from the IBA project. The blue line is a fitted straight line with the 95% confidence interval marked in gray. The equations of the fitted lines are shown in blue. **A:** The fitted line is forced through the origin, as the biomass is zero at zero individuals. The number of individuals per gram is estimated at 273+-17. The R-squared is 0.81. **B:** The same dataset with logged axes. The slope is 1.09+-0.07, that is, significantly larger than 1.0. This indicates that large samples (in terms of biomass) tend to have slightly more specimens in them than a strict proportional relationship between biomass and specimens would suggest. The R-squared is 0.79.

### Data records

The raw sequencing data (including the primer sequences) generated in this study are available at the European Nucleotide Archive (ENA) under study accession number PRJEB61109. Processed sequencing data include raw ASV sequences in FASTA format and ASV count files that contain the counts of each ASV (rows) in each sample (columns). It contains ASVs generated from Malaise trap samples (lysates, homogenates and preservative ethanol) and soil and litter samples. These data are organized by country (Sweden/Madagascar) and are available at https://doi.org/10.17044/scilifelab.25480681.v5 ^37^together with ‘sites_metadata’, ‘samples_metadata’, ‘sequencing_metadata’ and ‘spike_ins_metadata’ files. The same repository also contains datasets describing standing characteristics (‘stand_characteristics_MG.tsv’) and soil chemistry (‘soil_chemistry_SE.tsv’ and ‘soil_chemistry_MG.tsv’) collected at each Malaise trap location, as well as data on biomass and specimen counts (biomass_count_IBA.tsv and biomass_count_SIIP.tsv).

Processed ASV data include ASV files after taxonomic annotation with SINTAX and phylogenetic reassignment (‘asv_taxonomy.tsv’), sequences in FASTA format based on the representative ASV (see above) for each cluster (‘cluster_reps.fasta’), ASV cluster designations and taxonomy (‘cluster_taxonomy.tsv’), consensus taxonomy of clusters (‘cluster_consensus_taxonomy.tsv’) and summed ASV counts for each cluster (‘cluster_counts.tsv’). In addition, we provide taxonomy and count files for OTUs remaining after NEEAT filtering and cleaning to remove OTUs present in control samples. These files are available at https://doi.org/10.17044/scilifelab.27202368.v3 ^60^.

We also provide an earlier upload of raw ASV data with corresponding metadata files: https://doi.org/10.17044/scilifelab.25480681.v1 ^61^. This upload contains ASVs from Malaise trap samples processed with mild lysis only, with the exception of 15 samples for which we also provide data from homogenates and preservative ethanol. These data were processed using an earlier version of the bioinformatic processing workflow, and we also provide the data resulting from this effort ^62^. In this version, Swarm was run with setting ‘-d 13’, taxonomic re-assignment of ASVs using the insect phylogeny was not performed and removal of NUMTs and other potential noise was done using only taxonomic and abundance information. In summary, in this version of the workflow we removed ASVs present in more than 5% of blanks, ASVs unassigned at the family level, and clusters with less than 3 total counts summed across each dataset. For the Madagascar dataset, very few ASVs could be reliably assigned a taxonomy below the order level. Consequently, for the Madagascar dataset in this version we did not remove ASVs unassigned at the family level during the cleaning steps. In this version, homogenate samples from Sweden and soil and litter samples from both Sweden and Madagascar were not included, because they were unavailable at the time.

The data files provided contain ASVs that represent biological spike-ins. Specifically, the biological spike-ins used in Swedish datasets (lysates and homogenates) are *Gryllodes sigillatus* (Orthoptera: Gryllidae; in our datasets, the annotation of this taxon is *Gryllodes supplicans*; the reason is that this is the name used by GBIF for the matching CO1 sequences)*, Gryllus bimaculatus* (Orthoptera: Gryllidae), *Shelfordella lateralis* (Blattodea: Blattidae)*, Drosophila serrata, Drosophila bicornuta* and *Drosophila jambulina* (all Diptera: Drosophilidae; *D. jambulina* is annotated as ‘unclassified.Drosophila’ in our datasets due to name ambiguity in the annotation of the corresponding BIN in BOLD). The spike-ins used for Madagascar datasets (lysates) are *Orius majusculus* (Hemiptera: Anthocoridae), *Macrolophus pygmaeus* (Hemiptera: Miridae), *Delphastus pusillus* (Coleoptera: Coccinellidae), *Aphidius colemani* (Hymenoptera: Braconidae) *and Aphidoletes aphidimyza* (Diptera: Cecidomyiidae).

Homogenate samples (only available from Sweden) also contain synthetic spike-ins (raw homogenate data only). In the github repository https://github.com/insect-biome-atlas/utils, we provide tools to remove such ASVs and to use them to generate calibrated read numbers.

The ASVs corresponding to the OTUs that remained after NEEAT filtering and cleaning, together with their counts and metadata for the Swedish dataset, can also be accessed and viewed interactively through the ASV-portal (https://asv-portal.biodiversitydata.se)^63^ at the Swedish Biodiversity Infrastructure (SBDI), as well as through the Global Biodiversity Information Facility (GBIF) using https://doi.org/10.15468/veahzb for the lysate ^64^, https://doi.org/10.15468/af5cwp for the ethanol ^65^, https://doi.org/10.15468/awjycd for the homogenate ^66^, and https://doi.org/10.15468/783jyb for the soil-litter dataset ^67^. Here all ASVs (and their counts) are present rather than just OTU-representative ASVs.

### Technical Validation

Throughout all steps of sample processing we used negative control samples to control for cross-sample contamination. More specifically, we used negative controls during mild lysis and homogenization (‘buffer_blank’ and ‘buffer_blank_art_spikes’), DNA purification (‘extraction_neg’) and library preparation (‘pcr_neg’). Negative control samples can be identified in column ‘lab_sample_type’ in the ‘sequencing_metadata’ files available at https://doi.org/10.17044/scilifelab.25480681.v5 ^37^.

NUMTs and other types of noise were removed from the clustered ASV results using the NEEAT-filtering algorithm described above and we examined the median number of OTUs and sum of reads in negative controls and samples before and after this filtering step. For Swedish samples the median number of OTUs was 296 and 191, and the median sum of reads was 665k and 641k before and after filtering, respectively. For negative controls these numbers were 3 and 1 OTUs and 100 and 20 reads before and after filtering, respectively.

Samples in the Madagascar dataset had a median of 292 and 146 OTUs, and a median of 490k and 248k summed reads before and after filtering, respectively. Negative controls in this dataset had a median of 3 and 1 OTUs and 831 and 18 summed reads before and after filtering, respectively.

As part of data clean-up, it is usually advised to remove ASVs present in negative controls, or the maximum number of reads for those, from the entire dataset ^68^. However, after careful inspection of our negative controls, we noticed that only a few ASVs were persistently showing up in control samples. The majority of ASVs seemed to be arthropod sequences that were present in the bulk samples, and also sporadically present in negative controls in relatively small numbers. This was presumably due to DNA spreading between samples through tiny droplets during sample processing, or to low-level of “index hopping”, leading to incorrect assignment of reads during sequencing, despite the use of double-unique indexes in library preparation ^69^. Overall, this type of contamination was small and appeared random, leading us to the conclusion that removing all ASVs present in negative controls would impoverish the results without significantly improving quality. To ensure that we removed only true contaminants, we decided to filter out only those ASV clusters that showed up in more than 5% of negative controls that were successfully sequenced.

For Sweden, only seven clusters fit these criteria. Five of these were yeasts or bacteria; the remaining two were *Homo sapiens* (assigned to *H. neanderthalensis* due to GBIF removal of human data), and *Salticus scenicus* (zebra spider, a jumping spider). The latter is common in the building where samples were processed.

For Madagascar, 23 clusters fit the criteria. In addition to bacteria, yeasts, a plant (wheat or barley) and *Homo sapiens*, there were 14 arthropod clusters: *Allacma fusca*, *Entomobrya sp*., *Orchesella flavescens*, *O. cincta* and *Lepidocyrtus* sp. (Collembola); *Shelfordella lateralis* (Blattodea); *Nematopogon metaxella*, *Oligia* sp. (Lepidoptera); *Helina depuncta* and *Helina sp.*, *Leptogaster cylindrica*, *Sciapus platypterus*, *Sericomyia* sp. and *Sylvicola stackelbergi* (Diptera). Only *Entomobrya* sp. and *H. depuncta* occurred in more than 10% of control samples (13% and 12%, respectively). All of these clusters correspond to common species or spike-ins in the Swedish samples, which were processed before the Madagascar samples in the same lab. Many of them also occur in or around the building where samples were processed.

Some ASVs were assigned to a reference sequence in the BOLD database annotated as *Zoarces gillii* (BOLD:AEB5125), a fish found between Japan and eastern Korea. Closer inspection revealed that this was a mis-annotated bacterial sequence and ASVs assigned to this reference most likely represent bacterial sequences in our dataset. ASVs assigned to this species (232 ASVs in 55 OTUs for Sweden and 99 ASVs in 34 OTUs for Madagascar) were consequently removed. This record has been deleted from BOLD after our custom reference database was constructed.

After removal of OTUs present in >5% of negative controls, as well as the *Z. gillii* OTUs, the median number of OTUs and median sum of reads in negative controls for both Sweden and Madagascar was 0. Samples in Sweden had a median of 189 OTUs and 640k total read counts, respectively. For Madagascar, samples had a median of 145 OTUs and 248k total read counts (Table 2).

**Table 2.** Median number of OTUs and sum of read counts.

| Dataset | Output type | Sample type | OTUs | Sum of counts |
| --- | --- | --- | --- | --- |
| Sweden | Swarm | Sample | 296 | 665348 |
|  |  | Negative control | 3 | 100 |
|  | NEEAT-filtered | Sample | 191 | 640509 |
|  |  | Negative control | 1 | 20 |
|  | Cleaned | Sample | 189 | 640475 |
|  |  | Negative control | 0 | 0 |
| Madagascar | Swarm | Sample | 292 | 490458 |
|  |  | Negative control | 3 | 831 |
|  | NEEAT-filtered | Sample | 146 | 247892 |
|  |  | Negative control | 1 | 18 |
|  | Cleaned | Sample | 145 | 247892 |
|  |  | Negative control | 0 | 0 |

To assess whether the sampling and sequencing design was sufficient to characterize insect diversity at different levels, we performed three separate analyses. We performed analyses to assess (i) whether sequencing depth was sufficient to detect most species present in an individual sample (ii) whether the number of traps per site was sufficient to characterize the site-level community, (iii) whether the number of sites was sufficient to characterize the fauna of the region (i.e. the entirety of Sweden, or all forests of Madagascar). To answer these questions, we used functionality from the ‘vegan’ R package ^70^, specifically the ‘rarecurve’ (Figure 4a-b) and ‘specaccum’ (Figure 4c-f) functions.

**Figure 4.**
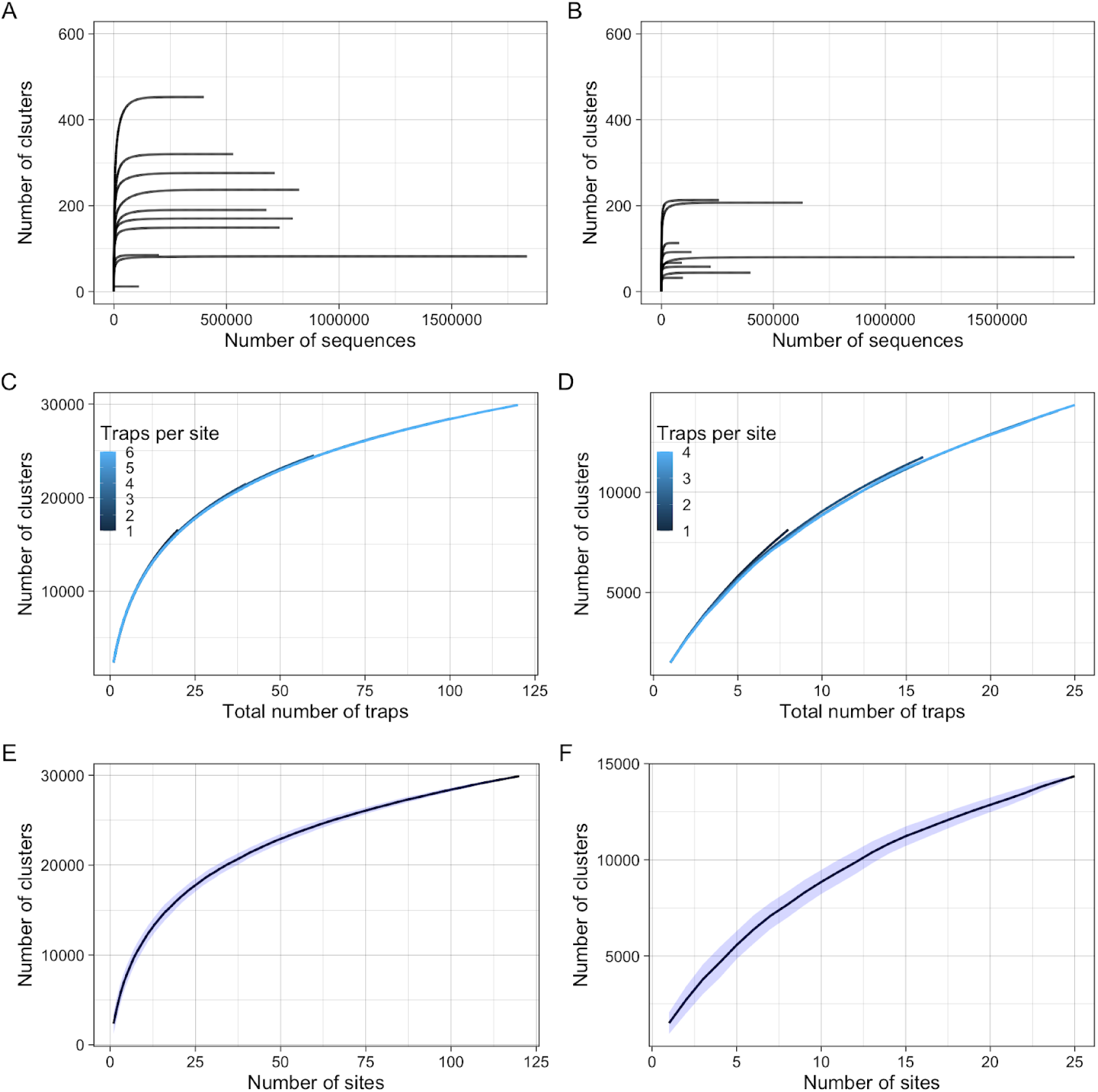
Results from the technical validation exercise to demonstrate sufficiency of the sequencing depth and the spatial sampling design to capture arthropod diversity in Sweden and Madagascar (left and right columns respectively. (A & B) Rarefaction curves illustrate cluster (species) accumulation with increasing sampling depth for 10 random samples in Sweden (A) and Madagascar (B). (C&D) Accumulation of species with an increasing number of traps for Sweden (C) and Madagascar (D). Each individual line represents the average accumulation of species given a specified site-level survey effort, that is, traps per site (indicated by colour). The X-axis (total number of traps), represents the pooled trap number across all multi-trap sites. (E&F) Accumulation of species with increasing spatial sampling effort (number of sites) for Sweden (E) and Madagascar (F).

To address the first question, we performed sample-level rarefaction on the sequencing depth of 10 random samples from both countries. (Figure 4a and Figure 4b). This allowed us to examine whether the number of sequences was sufficient to detect all species within a sample. As all curves level off before 500,000 reads, the average of 1.24 million reads per sample (median=1.09) was more than sufficient to detect the vast majority of species.

Within-site sampling efficiency was assessed using samples from sites that contained more than one trap (multitrap sites in Figure 1). To achieve this, we iteratively increased the number of traps representing each site and constructed species accumulation curves from samples pooled across all sites. For each iteration, we randomly sampled 20 permutations of *N* traps from each site (N = 1:6 traps per site for Sweden, N= 1:4 traps per site for Madagascar). For example, at the first iteration (N=1) a single permutation consisted of sampling one random trap per site - representing the lowest level of survey effort. In the second iteration (*N*=2), a single permutation consisted of two random traps per site. When *N* was equal to the maximum number of site-level traps (i.e. we were using the full sample), only one permutation was possible. For each permutation of traps, data was pooled and a species accumulation curve was constructed. Within each iteration (i.e. for each level of survey effort), the mean of the permutation-level curves were then taken to find the average species detection rate across sampling intensities. This analysis allows us to examine if, on average, more site-level sampling (i.e. traps) results in increased species detection. If lower within-site survey effort produces fewer species detections, we would expect lower intensities to produce less steep accumulation curves. The close proximity and trajectories of each species accumulation curve in panels 4C and 4D suggest that, on average, a single trap per location was generally sufficient to detect the same total number of species as higher sampling intensities.

Finally, at the country level, our analyses show that for Sweden, the total number of surveyed sites was bordering on sufficient to fully capture species richness across the country, as the curve begins to level off at higher sampling site numbers (Figure 4E). However, for Madagascar, the analysis suggests that 33 sites were still not sufficient to fully capture nation-wide species richness (Figure 4F).

In terms of taxonomic composition, the Malaise trap datasets are distinctly different from the soil and litter samples (Figure 5 and Figure 6). Malaise trap samples are dominated by Diptera and Hymenoptera in both countries, even though the Diptera are more prominent and the Hymenoptera less so in Madagascar. These orders are followed by the remaining three major insect orders (Coleoptera, Lepidoptera, and Hemiptera).

**Figure 5.**
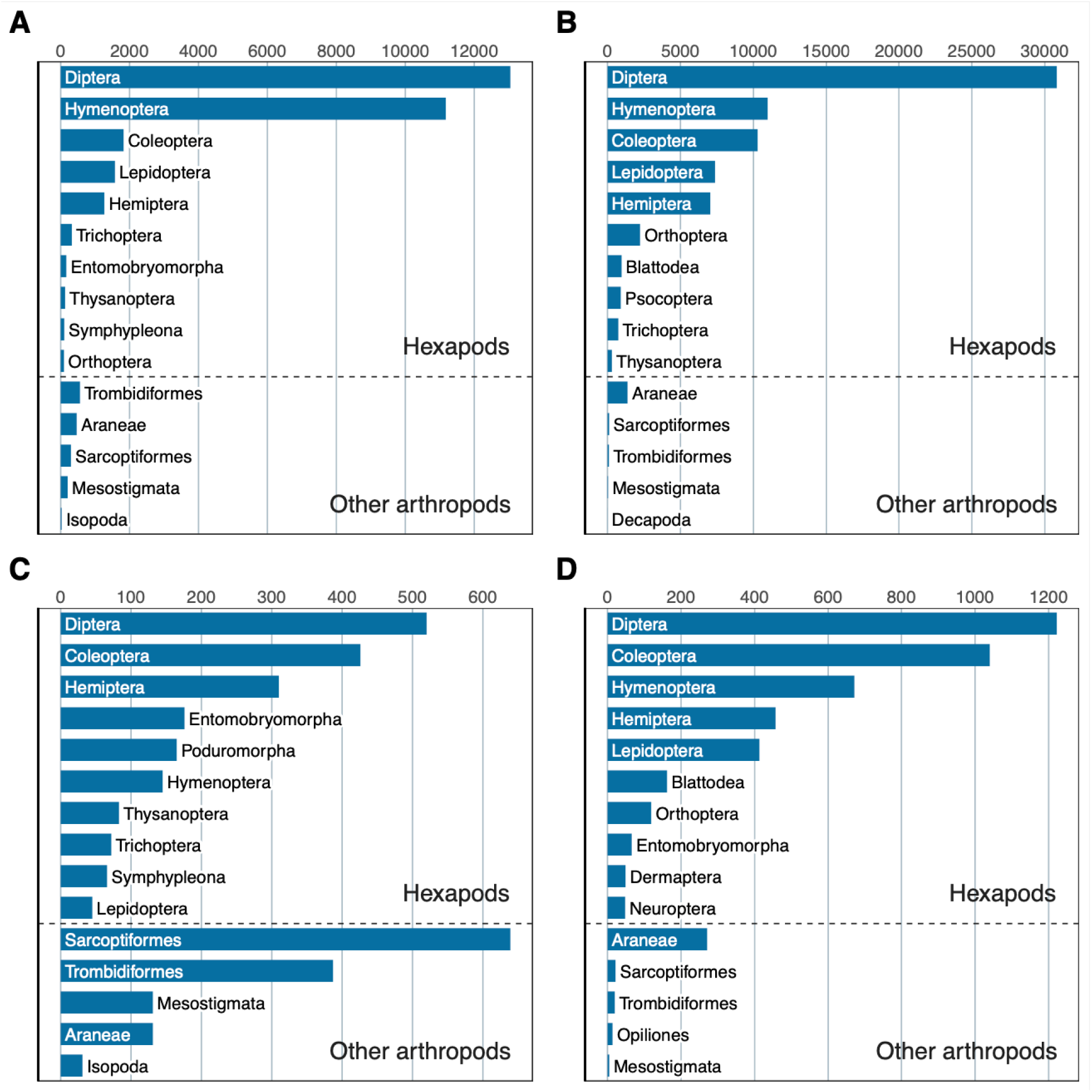
Order-level taxonomic composition of the IBA datasets (number of OTUs). (A) Swedish Malaise trap samples. (B) Malagasy Malaise trap samples. (C) Swedish soil and litter samples. (D) Malagasy litter samples. The bar plots show the top ten insect (Hexapoda) orders, and the top five other arthropod orders. Note that the Swedish datasets represent more intense sampling (four times as many samples) of a less diverse fauna than the Malagasy datasets. Also note that the Malagasy litter data does not include any soil samples, in contrast to the Swedish data.

**Figure 6.**
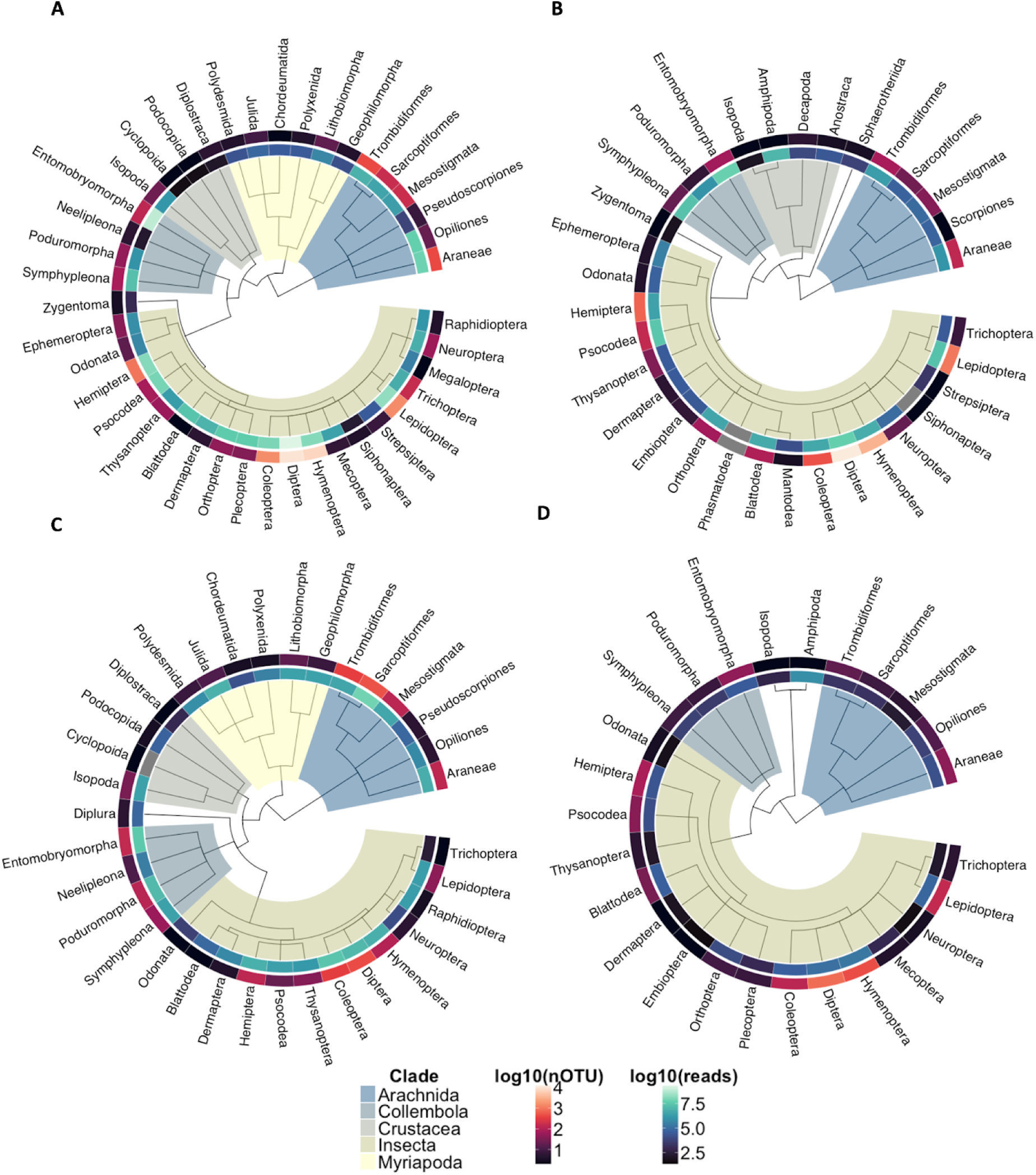
Order-level taxonomic composition of Swedish (A) and Malagasy (B) Malaise trap samples, Swedish soil and litter samples (C), Malagasy litter samples (D) and Swedish (C) and Malagasy (D) Malaise trap samples. Displayed are the Arthropod taxonomic trees with each tip representing an order. Major clades are highlighted across this tree. The inner ring around each order represents the base 10 logarithm of the total number of reads for each order and the outer ring represents the base 10 logarithm of the total number of assigned OTUs. For clarity, only orders with at least 100 reads are displayed.

Other arthropods constitute a relatively small proportion of the diversity. In the soil and litter samples, mites and spiders constitute a larger fraction of the diversity, particularly in Sweden. Among the insects, the bristletails (Entomobryomorpha and Poduromorpha) are also considerably more prominent than in Malaise trap samples. These patterns largely agree with expectations and suggest that the data properly reflects the diversity of the sampled faunas ^8,22,23,71^.

The precision and accuracy of the Swedish data are evaluated in ^31^as part of the methods development effort, and in more detail for Swedish Lepidoptera (butterflies and moths) in ^72^. Thanks to a long tradition of naturalists and the relative completeness of the reference libraries, the Swedish insect fauna permits powerful tests of the quality of the data from the Insect Biome Atlas project.

The 1,705 Lepidoptera OTUs found in Swedish Malaise trap samples (using version 1 of the data processing pipeline on lysates only) matched 1,535 unique species previously known from Sweden and included a number of tentative new species records ^72^. This suggests an over-splitting rate of less than 10%. The data covered 51% of the species ever recorded from the country, and 57% of the permanent resident species; the coverage was roughly the same across Lepidoptera families. Full-length barcoding of individual specimens representing 10 of the clusters that did not match known Swedish species confirmed that eight of them represented new species for the country or to science, or new and previously unknown CO1 variants at genetic distances that typically signal unique species.

Similar analyses across all families of Swedish insects also indicate good accuracy and precision of the processed IBA data ^31^. After chimera removal, clustering with swarm had a precision of 0.98 and a recall of 0.91 when assessed against the species annotation of the ASVs. That is, the clusters were highly consistent with taxonomy, but there was a slight tendency towards over-splitting as shown by the lower recall values. A considerable amount of this over-splitting disappeared after noise removal with the NEEAT algorithm. In the cleaned data, the vast majority of well-known Swedish insect families are represented by fewer clusters than described species, with only a few instances apparently representing substantial over-splitting. Many families are represented by slightly more than half of the recorded number of species, suggesting that the quality of the data is similar to that of Lepidoptera. The over-splitting of Orthoptera, an order notorious for its rampant NUMTs, is considerably lower in the IBA data than in comparable datasets ^73^.

### Usage Notes

The primers used for amplification were not removed from raw sequencing data and are therefore part of the reads deposited at the European Nucleotide Archive (ENA). The biological and synthetic spike-in sequences are also part of the raw data deposited at ENA. The clusters corresponding to spike-ins were not removed during the bioinformatic processing, so they are also included in the cleaned cluster data. The relationships between the metadata and ASV files are visualized in the simplified data model shown in Figure 7. The sample names used in the ASV and clustered files refer to the ‘sampleID_NGI’ field in the corresponding ‘sequencing_metadata’ file for each country (‘CO1_sequencing_metadata_SE.tsv’ and ‘CO1_sequencing_metadata_MG.tsv’). To retrieve metadata information about the original sample from which ASV data were obtained, you need to use the corresponding ‘sequencing_metadata’ file to identify the ‘sampleID_FIELD’ associated with the ‘sampleID_NGI’ that the ASV data refers to.

**Figure 7.**
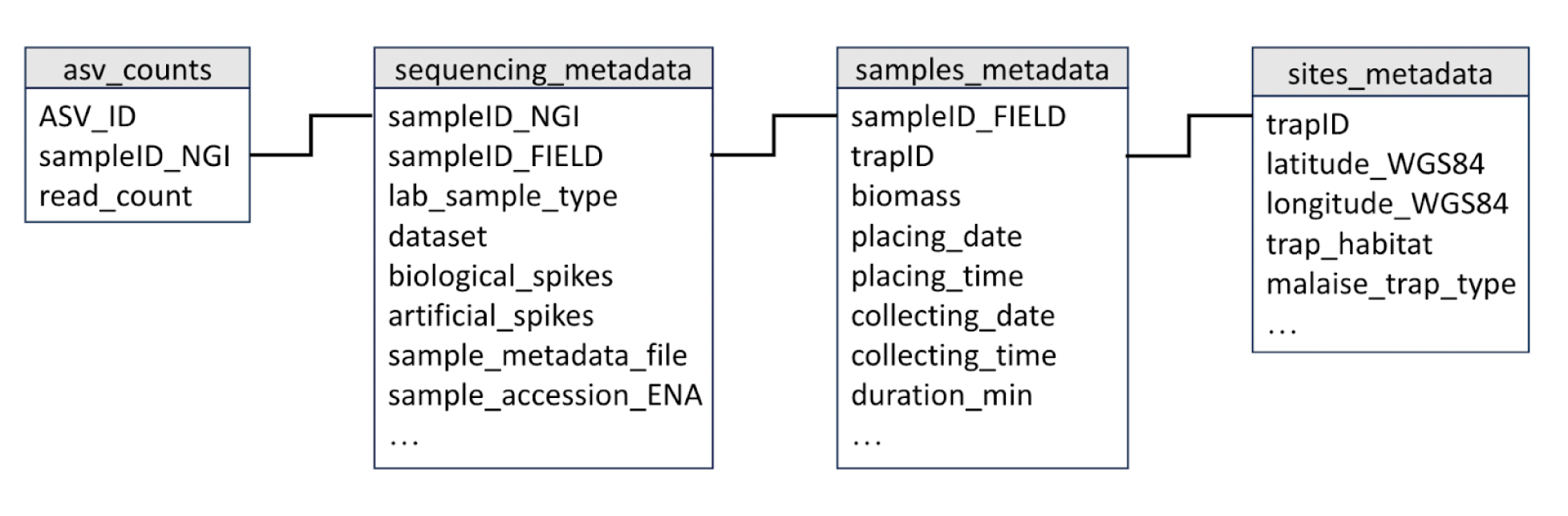
Simplified data model describing how the key fields in the main data and metadata files are linked.

Once you have the ‘sampleID_FIELD’ information, you can then use column ‘sample_metadata_file’ to identify the ‘samples_metadata’ file where all metadata associated with that specific sample (e.g. ‘placing_date’, ‘placing_time’, ‘biomass’, ‘trapID’, etc) is stored. From the ‘sites_metadata’ files you can retrieve all information associated with the sampling sites (e.g. ‘latitudeWGS84’, ‘longitudeWGS84’, ‘trap_habitat’). To identify the location of each sample, use the “trapID” column from the corresponding ‘sample_metadata’ file.

To facilitate usage of the data, we provide a set of scripts to process the data further, including scripts to remove spike-in reads and control samples, to use spike-in data (biological and synthetic) and weight data to generate calibrated read numbers and to generate distribution maps for Sweden or Madagascar based on site occurrence data. These scripts are available at https://github.com/insect-biome-atlas/utils.

### Other complementary datasets

At each sampling site in Sweden and Madagascar we collected information on ecosystem functions (including seed dispersal, predation, herbivory, pollination and decomposition) and arthropod biomass. These data are described and made available in ^32,32^. At each Malaise trap location in Sweden we also collected soil samples for biotic characterization (∼460 bp portion of the bacterial 16S rRNA gene, V3-V4 hypervariable region and ∼220-500 bp of the nuclear internal transcribed spacer 2 (ITS2) region of fungi). Finally, to characterize the microbiome associated with the insect community samples in Sweden, we amplified a ∼253 bp portion of the bacterial 16S rRNA gene (V4 hypervariable region) from the DNA extracted from the Malaise trap homogenates. These data will be published at a later date.

In Sweden, each Malaise trap was associated with a sampling plot from the National Inventory of Landscapes in Sweden (NILS) for which extensive data on land cover, land use and landscape structure spanning 18 years (2003-2020) are available (https://landskap.slu.se/nils/dv). NILS sampling square (‘NILS_square’) and sampling plot (‘NILS_plot’) associated with each IBA sampling location can be retrieved from ‘sites_metadata_SE.tsv’ file available at https://doi.org/10.17044/scilifelab.25480681.v561.

## Code Availability

Code for generating the statistics and plotting the figures presented in this paper are available on GitHub (https://github.com/insect-biome-atlas/paper-data/code).

## Acknowledgements

This work was supported by the Knut and Alice Wallenberg Foundation (grant KAW 2017.088 to FR), the Swedish Research Council (grant 2018-04620 to FR, grant 2021-03784 to AJMT, grant 2021-05563 to AFA), the Polish National Agency for Academic Exchange (grant PPN/PPO/2018/1/00015 to PL), the Polish National Science Centre (grant 2018/31/B/NZ8/01158 to PL), the European Research Council (Synergy Grant 856506 to TR), the Swedish University of Agricultural Sciences (Career Support grant to TR), the Science for Life Laboratory (SciLifeLab) & Wallenberg Data Driven Life Science Program funded by Knut and Alice Wallenberg Foundation (grants: KAW 2020.0239 and KAW 2017.0003), the National Bioinformatics Infrastructure Sweden (NBIS) at SciLifeLab, the Abisko Scientific Research Station and the Swedish Infrastructure for Ecosystem Science (Swedish Research Council grant no. 4.3-2021-00164) and the strategic research area BECC - Biodiversity and Ecosystem services in a Changing Climate. The computations and data handling were enabled by resources in projects SNIC 2020/15-307, SNIC 2022/5-211, NAISS 2023/5-209, and NAISS 2024/5-207 (compute) and SNIC 2020/16-248 (storage), provided by the National Academic Infrastructure for Supercomputing in Sweden (NAISS) and the Swedish National Infrastructure for Computing (SNIC) at Uppsala Multidisciplinary Center for Advanced Computational Science (UPPMAX), partially funded by the Swedish Research Council (grant 2022-06725 and 2018-05973). The authors also acknowledge support from the National Genomics Infrastructure in Stockholm and from the Swedish Biodiversity Data Infrastructure (SBDI). For the work carried out in Madagascar, the authors acknowledge support from the Madagascar National Parks, Fanamby, Asity, Groupe d’Etudes et de Recherches sur les Primates (GERP), Centre National de Formation, d’Etudes et de Recherches en Environnement et Foresterie (CNFEREF), Missouri Botanical Garden (MBG), Ambatovy, Ministère de l’Environnement et du Développement Durable, Parc Botanique et Zoologique de Tsimbazaza and Laboratoires des Radioisotopes. Finally we would like to thank Paula Arribas for helping with the protocol for DNA extraction of soil and leaf litter arthropod communities.

## Author contributions

**Andreia Miraldo**: conceptualization, data curation, formal analysis, investigation, methodology, project administration, supervision, validation, writing-original draft, writing-review and editing; **John Sundh:** conceptualization, data curation, formal analysis, investigation, methodology, validation, writing-original draft, writing-review and editing; **Elzbieta Iwaszkiewicz-Eggebrecht:** investigation, methodology, writing-review and editing; **Mateusz Buczek:** investigation, methodology, writing-review and editing; **Robert Goodsell:** investigation, writing-review and editing; **Håkan Johansson:** investigation, writing-review and editing; **Brian L. Fisher:** investigation, resources, writing-review and editing; **Dimby Raharinjanahary:** investigation, writing-review and editing; **Eric T. Rajoelison:** investigation, writing-review and editing; **Chrislain Ranaivo:** investigation, writing-review and editing; **Claver E. Randrianandrasana:** investigation, writing-review and editing; **Jean-Jaques Rafanomezantsoa:** investigation, writing-review and editing; **Lokeshwaran Manoharan:** investigation, writing-review and editing; **Emma Granqvist:** investigation, formal analysis, writing-review and editing; **Laura J. A. van Dijk:** investigation, writing-review and editing; **Louise Ahlberg:** investigation, writing-review and editing; **David Åhlén:** investigation, writing-review and editing; **Maj Aspebo:** investigation, writing-review and editing; **Staffan Åström**: investigation, writing-review and editing; **Anna Bellviken:** investigation, writing-review and editing; **Per-Erik Bergman:** investigation, writing-review and editing; **Sofie Björklund:** investigation, writing-review and editing; **Mats P. Björkman:** investigation, writing-review and editing; **Junchen Deng:** investigation, writing-review and editing; **Luke Desborough:** investigation, writing-review and editing; **Elisabet Dolff:** investigation, writing-review and editing; **Angelika Eliasson:** investigation, writing-review and editing; **Helena Elmquist:** investigation, writing-review and editing; **Hanna Z. S. Emanuelsson:** investigation, writing-review and editing; **Rebecka Erixon:** investigation, writing-review and editing; **Lennart Fahlen:** investigation, writing-review and editing; **Carina Frogner:** investigation, writing-review and editing; **Pieter Fürst:** investigation, writing-review and editing; **Andreas Grabs:** investigation, writing-review and editing; **Håkan Grudd:** investigation, writing-review and editing; **Daniela Guasconi:** investigation, writing-review and editing; **Marie Gunnarsson:** investigation, writing-review and editing; **Sibylle Häggqvist:** investigation, writing-review and editing; **Annika Hed:** investigation, writing-review and editing; **Eva Hörnström:** investigation, writing-review and editing; **Helene Johansson:** investigation, writing-review and editing; **Anna Jönsson:** investigation, writing-review and editing; **Sara Kanerot:** investigation, writing-review and editing; **Andreas Karlsson:** investigation, writing-review and editing; **Dave Karlsson:** investigation, writing-review and editing; **Mårten J. Klinth:** investigation, writing-review and editing; **Thomas Kraft:** investigation, writing-review and editing; **Rolf Lahti:** investigation, writing-review and editing; **Malin M. Larsson:** investigation, writing-review and editing; **Håkan Lernefalk:** investigation, writing-review and editing; **Ylva Lestander:** investigation, writing-review and editing; **Lars-Ture Lindholm:** investigation, writing-review and editing; **Märith Lindholm:** investigation, writing-review and editing; **Ulrika Ljung:** investigation, writing-review and editing; **Krister Ljung:** investigation, writing-review and editing; **Johannes Lundberg:** investigation, writing-review and editing; **Erik J. L. Lundin:** investigation, writing-review and editing; **Marie Malmenius:** investigation, writing-review and editing; **Daniel Marquina:** investigation, writing-review and editing; **Johan Martinelli:** investigation, writing-review and editing; **Laurent Mertz:** investigation, writing-review and editing; **Jan Nilsson:** investigation, writing-review and editing; **Aurora Patchett:** investigation, writing-review and editing; **Nannie L. Persson:** investigation, writing-review and editing; **John Persson:** investigation, writing-review and editing; **Monika Prus-Frankowska:** investigation, writing-review and editing; **Emilia D. E. Regazzoni:** investigation, writing-review and editing; **Karl-Gunnar Rosander:** investigation, writing-review and editing; **Mats Rydgård:** investigation, writing-review and editing; **Christina Sandblom:** investigation, writing-review and editing; **Jimmy Skord:** investigation, writing-review and editing; **Thomas Stålhandske:** investigation, writing-review and editing; **Fredric Svensson:** investigation, writing-review and editing; **Scarlett Szpryngiel:** investigation, writing-review and editing; **Kristina Tajani:** investigation, writing-review and editing; **Maud I. Tyboni:** investigation, writing-review and editing; **Carina R. Ugarph:** investigation, writing-review and editing; **Leif Vestermark:** investigation, writing-review and editing; **Jakob Vilhelmsson:** investigation, writing-review and editing; **Niclas Wahlgren:** investigation, writing-review and editing; **Anett Wass:** investigation, writing-review and editing; **Per Wetterstrand:** investigation, writing-review and editing; **Piotr Łukasik:** conceptualization, resources, supervision, writing-original draft, writing-review and editing; **Ayco J. M. Tack:** conceptualization, funding acquisition, resources, supervision, writing-original draft, writing-review and editing; **Anders F. Andersson:** conceptualization, funding acquisition, resources, supervision, writing-original draft, writing-review and editing; **Tomas Roslin**: conceptualization, funding acquisition, resources, supervision, writing-original draft, writing-review and editing; **Fredrik Ronquist:** conceptualization, funding acquisition, resources, supervision, validation, writing-original draft, writing-review and editing.

## References

1. Sverdrup-Thygeson A. Extraordinary Insects: Weird, Wonderful, Indispensable: The Ones Who Run the World. UK edition. Mudlark; 2019.

2. Crespo-Pérez V, Kazakou E, Roubik DW, Cárdenas RE. The importance of insects on land and in water: a tropical view. Vectors Med Vet Entomol • Spec Sect Insects UN Sustain Dev Goals. 2020;40:31–38. doi:10.1016/j.cois.2020.05.016

3. Grames EM, Montgomery GA, Youngflesh C, Tingley MW, Elphick CS. The effect of insect food availability on songbird reproductive success and chick body condition: Evidence from a systematic review and meta-analysis. Ecol Lett. 2023;26(4):658–673. doi:10.1111/ele.14178

4. Kagata H, Ohgushi T. Bottom-up trophic cascades and material transfer in terrestrial food webs. Ecol Res. 2006;21(1):26–34. doi:10.1007/s11284-005-0124-z

5. Yang LH, Gratton C. Insects as drivers of ecosystem processes. Ecology. 2014;2:26–32. doi:10.1016/j.cois.2014.06.004

6. Mora C, Tittensor DP, Adl S, Simpson AGB, Worm B. How Many Species Are There on Earth and in the Ocean? PLoS Biol. 2011;9(8):e1001127. doi:10.1371/journal.pbio.1001127

7. Stork NE. How Many Species of Insects and Other Terrestrial Arthropods Are There on Earth? Annu Rev Entomol. 2018;63(1):31–45. doi:10.1146/annurev-ento-020117-043348

8. Srivathsan A, Ang Y, Heraty JM, et al. Convergence of dominance and neglect in flying insect diversity. Nat Ecol Evol. 2023;7(7):1012–1021. doi:10.1038/s41559-023-02066-0

9. Hallmann CA, Sorg M, Jongejans E, et al. More than 75 percent decline over 27 years in total flying insect biomass in protected areas. PLOS ONE. 2017;12(10):e0185809. doi:10.1371/journal.pone.0185809

10. Lister BC, Garcia A. Climate-driven declines in arthropod abundance restructure a rainforest food web. Proc Natl Acad Sci. 2018;115(44):E10397–E10406. doi:10.1073/pnas.1722477115

11. Møller AP. Parallel declines in abundance of insects and insectivorous birds in Denmark over 22 years. Ecol Evol. 2019;9(11):6581–6587. doi:10.1002/ece3.5236

12. Outhwaite CL, McCann P, Newbold T. Agriculture and climate change are reshaping insect biodiversity worldwide. Nature. 2022;605(7908):97-102. doi:10.1038/s41586-022-04644-x

13. Seibold S, Gossner MM, Simons NK, et al. Arthropod decline in grasslands and forests is associated with landscape-level drivers. Nature. 2019;574(7780):671-674. doi:10.1038/s41586-019-1684-3

14. Thomas CD, Jones TH, Hartley SE. “Insectageddon”: A call for more robust data and rigorous analyses. Glob Change Biol. 2019;25(6):1891–1892. doi:10.1111/gcb.14608

15. van Klink R, Bowler DE, Gongalsky KB, Shen M, Swengel SR, Chase JM. Disproportionate declines of formerly abundant species underlie insect loss. Nature. Published online December 20, 2023:1–6. doi:10.1038/s41586-023-06861-4

16. van Klink R, Bowler DE, Gongalsky KB, Swengel AB, Gentile A, Chase JM. Meta-analysis reveals declines in terrestrial but increases in freshwater insect abundances. Science. 2020;368(6489):417-420. doi:10.1126/science.aax9931

17. Basset Y, Lamarre GPA. Toward a world that values insects. Science. 2019;364(6447):1230-1231. doi:10.1126/science.aaw7071

18. Cardoso P, Barton PS, Birkhofer K, et al. Scientists’ warning to humanity on insect extinctions. Biol Conserv. 2020;242:108426. doi:10.1016/j.biocon.2020.108426

19. Harvey JA, Tougeron K, Gols R, et al. Scientists’ warning on climate change and insects. Ecol Monogr. 2023;93(1):e1553. doi:10.1002/ecm.1553

20. Harvey JA, Heinen R, Armbrecht I, et al. International scientists formulate a roadmap for insect conservation and recovery. Nat Ecol Evol. 2020;4(2):174–176. doi:10.1038/s41559-019-1079-8

21. Wilson RJ, Fox R. Insect responses to global change offer signposts for biodiversity and conservation. Ecol Entomol. 2021;46(4):699–717. doi:10.1111/een.12970

22. Karlsson D, Hartop E, Forshage M, Jaschhof M, Ronquist F. The Swedish Malaise Trap Project: A 15 Year Retrospective on a Countrywide Insect Inventory. Biodivers Data J. 2020;8:e47255. doi:10.3897/BDJ.8.e47255

23. Ronquist F, Forshage M, Häggqvist S, et al. Completing Linnaeus’s inventory of the Swedish insect fauna: Only 5,000 species left? PLOS ONE. 2020;15(3):e0228561. doi:10.1371/journal.pone.0228561

24. van Klink R, August T, Bas Y, et al. Emerging technologies revolutionise insect ecology and monitoring. Trends Ecol Evol. 2022;37(10):872–885. doi:10.1016/j.tree.2022.06.001

25. Hewitt GM. Post-glacial re-colonization of European biota. Biol J Linn Soc. 1999;68(1):87–112. doi:10.1006/bijl.1999.0332

26. Antonelli A, Smith RJ, Perrigo AL, et al. Madagascar’s extraordinary biodiversity: Evolution, distribution, and use. Science. 2022;378(6623):eabf0869. doi:10.1126/science.abf0869

27. Ståhl G, Allard A, Esseen PA, et al. National Inventory of Landscapes in Sweden (NILS)—scope, design, and experiences from establishing a multiscale biodiversity monitoring system. Environ Monit Assess. 2011;173(1):579–595. doi:10.1007/s10661-010-1406-7

28. Iwaszkiewicz-Eggebrecht E, Łukasik P, Buczek M, et al. FAVIS: Fast and versatile protocol for non-destructive metabarcoding of bulk insect samples. PloS One. 2023;18(7):e0286272. doi:10.1371/journal.pone.0286272

29. Iwaszkiewicz-Eggebrecht E, Łukasik P, Buczek M, et al. FAVIS: Fast and Versatile protocol for metabarcoding of bulk Insect Samples from large-scale diversity monitoring projects v2. protocols.io. Published online November 22, 2023. doi:10.17504/protocols.io.kqdg36261g25/v2

30. Persson N, Johansson H, Iwaszkiewicz-Eggebrecht E, Miraldo A. Sample homogenization and DNA extraction for bulk insect catches. protocols.io. Published online 2024. doi:10.17504/protocols.io.yxmvm31bbl3p/v1

31. Sundh J, Granqvist E, Iwaszkiewicz-Eggebrecht E, et al. HAPP: High-Accuracy Pipeline for Processing of deep metabarcoding data. Published online 2024.

32. van Dijk LJA, Miraldo A, Raharinjanahary D, et al. Biotic and abiotic drivers of ecosystem functioning differ between a temperate and a tropical region. Published online 2024. 10.1101/2024.02.28.582312

33. van Dijk LJA, Fisher BL, Miraldo A, et al. Temperature and water availability drive insect seasonality across a temperate and a tropical region. Proceedings of the Royal Society B. 291(20240090). 10.1098/rspb.2024.0090

34. Ratnasingham S, Hebert PDN. bold: The Barcode of Life Data System (http://www.barcodinglife.org). Mol Ecol Notes. 2007;7(3):355–364. doi:10.1111/j.1471-8286.2007.01678.x

35. Tullgren A. Ein sehr einfacher Ausleseapparat für terricole Tierformen. Z Für Angew Entomol. 1918;4(1):149–150. doi:10.1111/j.1439-0418.1918.tb00820.x

36. Besuchet C, Burckhardt DH, Löbl I. The “Winkler/Moczarski” Eclector as an Efficient Extractor for Fungus and Litter Coleoptera. The Coleopterists Bulletin. 1987;41(4):392–394.

37. Miraldo A, Iwaszkiewicz-Eggebrecht E, Sundh J, et al. Amplicon sequence variants from the Insect Biome Atlas project. Published online 2024. 10.17044/scilifelab.25480681.v5

38. Iwaszkiewicz-Eggebrecht E, Prus-Frankowska M, Łukasik P. Synthetic COI spike-ins for use in metabarcoding-based insect biodiversity surveys. protocols.io. Published online 2023. doi:10.17504/protocols.io.14egn33ryl5d/v2

39. Arribas P, Andújar C, Salces-Castellano A, Emerson BC, Vogler AP. The limited spatial scale of dispersal in soil arthropods revealed with whole-community haplotype-level metabarcoding. Mol Ecol. 2021;30(1):48–61. doi:10.1111/mec.15591

40. Arribas P, Andújar C, Hopkins K, Shepherd M, Vogler AP. Metabarcoding and mitochondrial metagenomics of endogean arthropods to unveil the mesofauna of the soil. Methods Ecol Evol. 2016;7(9):1071–1081. doi:10.1111/2041-210X.12557

41. Glenn TC, Nilsen RA, Kieran TJ, et al. Adapterama I: universal stubs and primers for 384 unique dual-indexed or 147,456 combinatorially-indexed Illumina libraries (iTru & iNext). Lazo G, ed. PeerJ. 2019;7:e7755. doi:10.7717/peerj.7755

42. Elbrecht V, Braukmann TWA, Ivanova NV, et al. Validation of COI metabarcoding primers for terrestrial arthropods. Pochon X, ed. PeerJ. 2019;7:e7745. doi:10.7717/peerj.7745

43. Elbrecht V, Leese F. Validation and Development of COI Metabarcoding Primers for Freshwater Macroinvertebrate Bioassessment. Front Environ Sci. 2017;5. https://www.frontiersin.org/articles/10.3389/fenvs.2017.00011

44. Bonath F. Increased Complexity of Amplicon Libraries Using Phased Primers. National Genomics Infrastructure; 2021.

45. Wu L, Wen C, Qin Y, et al. Phasing amplicon sequencing on Illumina Miseq for robust environmental microbial community analysis. BMC Microbiol. 2015;15(1):125. doi:10.1186/s12866-015-0450-4

46. Glenn TC, Pierson TW, Bayona-Vásquez NJ, et al. Adapterama II: universal amplicon sequencing on Illumina platforms (TaggiMatrix). PeerJ. 2019;7:e7786. doi:10.7717/peerj.7786

47. Martin M. Cutadapt removes adapter sequences from high-throughput sequencing reads. EMBnetjournal Vol 17 No 1 Gener Seq Data Anal. Published online 2011. doi:10.14806/ej.17.1.200

48. Straub D, Blackwell N, Langarica-Fuentes A, Peltzer A, Nahnsen S, Kleindienst S. Interpretations of Environmental Microbial Community Studies Are Biased by the Selected 16S rRNA (Gene) Amplicon Sequencing Pipeline. Front Microbiol. 2020;11. https://www.frontiersin.org/journals/microbiology/articles/10.3389/fmicb.2020.550420

49. Callahan BJ, McMurdie PJ, Rosen MJ, Han AW, Johnson AJA, Holmes SP. DADA2: High-resolution sample inference from Illumina amplicon data. Nat Methods. 2016;13(7):581–583. doi:10.1038/nmeth.3869

50. Bohmann K, Elbrecht V, Carøe C, et al. Strategies for sample labelling and library preparation in DNA metabarcoding studies. Mol Ecol Resour. 2022;22(4):1231–1246. doi:10.1111/1755-0998.13512

51. Kircher M, Sawyer S, Meyer M. Double indexing overcomes inaccuracies in multiplex sequencing on the Illumina platform. Nucleic Acids Res. 2012;40(1):e3–e3. doi:10.1093/nar/gkr771

52. Edgar RC. SINTAX: a simple non-Bayesian taxonomy classifier for 16S and ITS sequences. bioRxiv. Published online January 1, 2016:074161. doi:10.1101/074161

53. Rognes T, Flouri T, Nichols B, Quince C, Mahé F. VSEARCH: a versatile open source tool for metagenomics. Hrbek T, ed. PeerJ. 2016;4:e2584. doi:10.7717/peerj.2584

54. Sundh J. COI reference sequences from BOLD DB. SciLifeLab. Published online 2022. 10.17044/scilifelab.20514192.v4

55. Barbera P, Kozlov AM, Czech L, et al. EPA-ng: Massively Parallel Evolutionary Placement of Genetic Sequences. Syst Biol. 2019;68(2):365–369. doi:10.1093/sysbio/syy054

56. Chesters D. Construction of a Species-Level Tree of Life for the Insects and Utility in Taxonomic Profiling. Syst Biol. 2017;66(3):426–439. doi:10.1093/sysbio/syw099

57. Czech L, Barbera P, Stamatakis A. Genesis and Gappa: processing, analyzing and visualizing phylogenetic (placement) data. Bioinformatics. 2020;36(10):3263–3265. doi:10.1093/bioinformatics/btaa070

58. Edgar RC, Haas BJ, Clemente JC, Quince C, Knight R. UCHIME improves sensitivity and speed of chimera detection. Bioinformatics. 2011;27(16):2194–2200. doi:10.1093/bioinformatics/btr381

59. Mahé F, Rognes T, Quince C, de Vargas C, Dunthorn M. Swarm: robust and fast clustering method for amplicon-based studies. Cohan F, ed. PeerJ. 2014;2:e593. doi:10.7717/peerj.593

60. Miraldo A, Iwaszkiewicz-Eggebrecht E, Sundh J, et al. Processed ASV data from the Insect Biome Atlas Project. Published online 2024. 10.17044/scilifelab.27202368.v3

61. Miraldo A, Iwaszkiewicz-Eggebrecht E, Sundh J, et al. Amplicon sequence variants from the Insect Biome Atlas project. Published online 2024. 10.17044/scilifelab.25480681.v1

62. Miraldo A, Iwaszkiewicz-Eggebrecht E, Sundh J, et al. Processed ASV data from the Insect Biome Atlas Project. Published online 2024. 10.17044/scilifelab.27202368.v1

63. Prager M, Lundin D, Ronquist F, Andersson AF. ASV portal: an interface to DNA-based biodiversity data in the Living Atlas. BMC Bioinformatics. 2023;24(1):6. doi:10.1186/s12859-022-05120-z

64. Miraldo A, Iwaszkiewicz-Eggebrecht E, Sundh J, et al. CO1 Amplicon Sequence Variants of bulk arthropod samples (mild lysis) collected with Malaise traps from the Insect Biome Atlas project in Sweden. Published online 2024. 10.15468/veahzb

65. Miraldo A, Iwaszkiewicz-Eggebrecht E, Sundh J, et al. CO1 Amplicon Sequence Variants of bulk arthropod samples (preservative ethanol) collected with Malaise traps from the Insect Biome Atlas project in Sweden. Version 1.1. Published online 2024. 10.15468/af5cwp

66. Miraldo A, Iwaszkiewicz-Eggebrecht E, Sundh J, et al. CO1 Amplicon Sequence Variants of bulk arthropod samples (homogenized) collected with Malaise traps from the Insect Biome Atlas project in Sweden. Published online 2024. 10.15468/awjycd

67. Miraldo A, Iwaszkiewicz-Eggebrecht E, Sundh J, et al. CO1 Amplicon Sequence Variants of soil and leaf litter arthropod communities collected at Malaise traps from the Insect Biome Atlas project in Sweden. Version 1.1. Published online 2024. 10.15468/783jyb

68. Alberdi A, Aizpurua O, Gilbert MTP, Bohmann K. Scrutinizing key steps for reliable metabarcoding of environmental samples. Methods Ecol Evol. 2018;9(1):134–147. doi:10.1111/2041-210X.12849

69. MacConaill LE, Burns RT, Nag A, et al. Unique, dual-indexed sequencing adapters with UMIs effectively eliminate index cross-talk and significantly improve sensitivity of massively parallel sequencing. BMC Genomics. 2018;19(1):30. doi:10.1186/s12864-017-4428-5

70. Oksanen J, Simpson GL, Blanchet FG, et al. vegan: Community Ecology Package. Published online May 21, 2024. Accessed May 24, 2024. https://cran.r-project.org/web/packages/vegan/index.html

71. Coleman D, Callaham M, Jr C. Fundamentals of Soil Ecology: Third Edition.; 2017.

72. Iwaszkiewicz-Eggebrecht E, Goodsell RM, Bengsson BÅ, et al. High-throughput biodiversity surveying sheds new light on the brightest of insect taxa. bioRxiv. Published online January 1, 2024:2024.10.25.620209. doi:10.1101/2024.10.25.620209

73. Buchner D, Sinclair JS, Ayasse M, et al. Upscaling biodiversity monitoring: Metabarcoding estimates 31,846 insect species from Malaise traps across Germany. Mol Ecol Resour. 2024;n/a(n/a):e14023. doi:10.1111/1755-0998.14023

